# Evaluation of origin of driving force for loop formation in a chromatin fiber

**DOI:** 10.1101/2020.06.24.168757

**Authors:** Hiroshi Yokota, Masashi Tachikawa

**Author notes:** These authors contributed equally to this work. Membership list can be found in the Acknowledgments section.

## Abstract

Chromosome condensation results from the formation of consecutive chromatin loops in which excluded volume interactions lead to chromosome stiffness. Formation of chromatin loops requires energy, but the source of such energy remains controversial. Here, we quantified the energy balance during chromatin loop formation by calculating the free energies of unlooped and looped chromatins using a lattice model of polymer chains. We tested two hypothetical energy sources: thermal fluctuation and ATP hydrolysis. We evaluated the free energy difference of the chain loop model without accounting for excluded volume interactions (phantom loop model), and integrated those interactions by employing the mean-field theory (interacting loop model), where we introduced the parameter of excluded volume interaction within a single loop *v*_ex_. Using our strategy, we confirmed that loop-growth efficiency calculated by the phantom loop model is too high to explain the experimental data. Comparing loop-growth efficiencies for each energy source, and using the interacting loop model, we found that excluded volume interaction is essential for chromatin’s resistance to looping, regardless of the energy source. We predict that the quantitative measurement of *v*_ex_ determines which energy source is more plausible.

**Author summary:** Before mitosis, the chromatin fibers of eukaryotic cells fold into consecutive loop structures and condense into rod-like chromosomes. Chromosome stiffness results from the interaction of the excluded volume between chromatin loops. The driving force of loop formation and growth is still controversial, despite the many efforts undertaken to clarify it. Two possible origins can be considered: the energy provided by thermal fluctuations or the energy gained from ATP hydrolysis. To discuss the validity of each, we constructed a theoretical model of chromatin loop formation that includes excluded volume interactions. Using this model, we calculated the free energy difference before and after chromatin loop formation, which corresponds to the energy that fuels chromatin looping. By comparing the results for each energy source, we conclude that the spatial distribution of chromatin loops should be relatively wide, given the large excluded volume interaction within a single loop, irrespective of which energy source is valid. Moreover, our results imply that intra-loop interactions are key to determine the driving force of chromatin loop formation.

## Introduction

Prior to mitosis, the chromatin fibers of eukaryotic cells fold into consecutive loop structures and condense into rod-like chromosomes (see Fig. 1 (a)). The stiffness of chromosomes results from the interaction of the excluded volume between each chromatin loop. One of the essential molecules for chromatin loop formation is a five-subunit protein complex named condensin, which belongs to the highly conserved family of SMC complexes.

**Fig 1.**
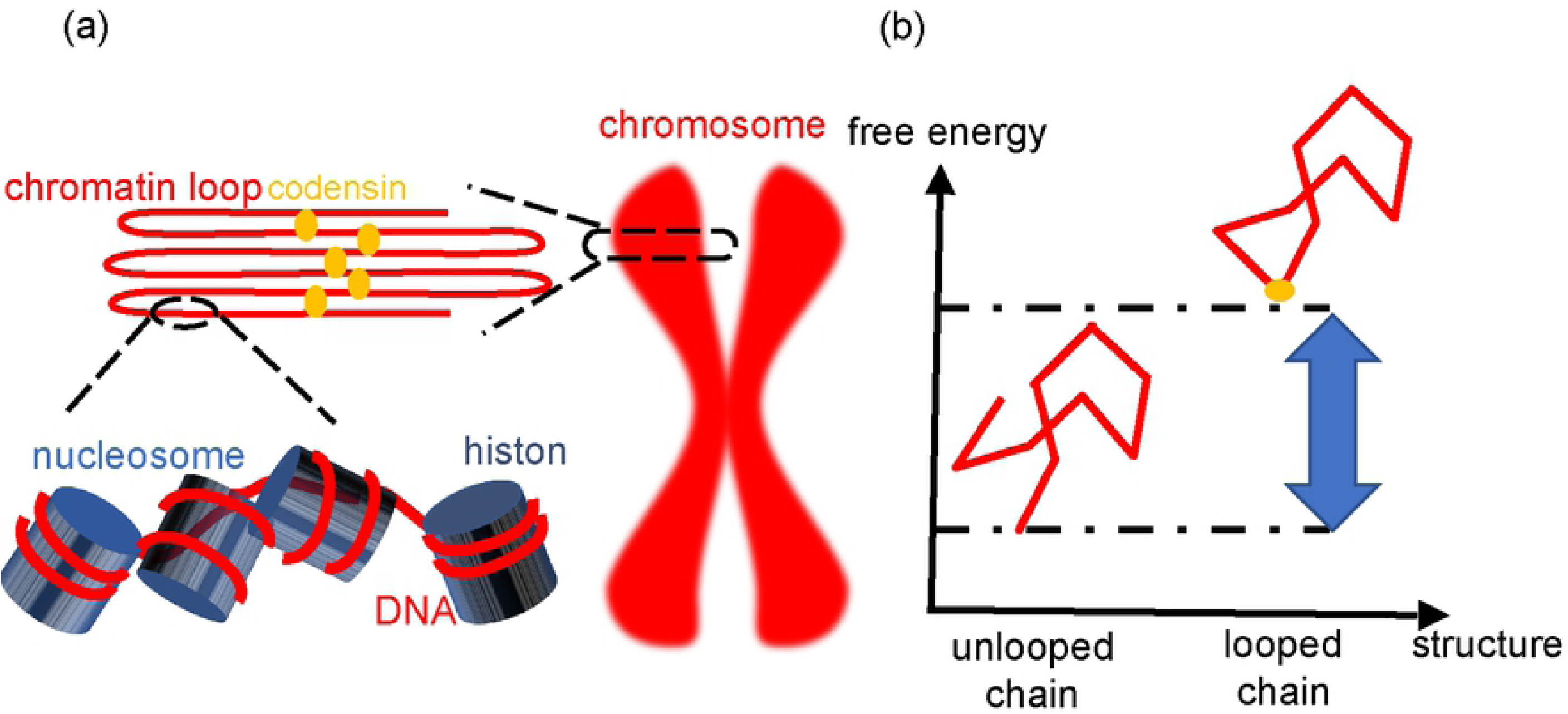
(a) Schematic picture of the chromosome composed of the consecutive chromatin loops. (b) The methodology which we used in this study. The energy source of the driving force for creating a loop is quantified by using the free energy difference before and after chromatin loop (s) formation.

Chromosome assembly is an attractive topic of study, both for theoretical [1–3] and experimental [4–6] researches. The mechanism and detailed dynamics of loop formation have been extensively studied yet they remain incompletely understood. One of the most promising hypotheses is that of loop extrusion, proposed by Alipour et al. [2]. In this hypothesis condensin binds at two neighboring sites in the chromatin fiber, and extrudes (pushes) it to form and enlarge a DNA loop. Loop extrusion has been theoretically modeled based on stochastic [7] and coarse-grained molecular dynamics simulations [1, 3]. Moreover, the loop extrusion activity of condensin on bare DNA was observed experimentally by Ganji et al. [6].

The driving force of loop formation and growth is also an important topic and a matter of debate. In general, two possible driving forces are considered. One is the direct power stroke of condensin as a motor protein coupled with its ATP hydrolysis. Recent experiments revealed the ATP-dependent translocation and loop formation activity of condensins along DNA [4, 6] and Terakawa et al. [4] proposed some motor activity models of condensins. The other driving force candidate is thermal fluctuation. The model proposed by Marko et al. [8] outlines how thermal fluctuation can act as the driving force for loop formation and growth. According to this model, the DNA fiber and condensin undergo a cyclic reaction involving conformational changes in condensin and several DNA forms captured by condensin. During this reaction, condensin binds to DNA and additionally captures a loop that was stochastically formed nearby. Then, it releases the former DNA binding site and rebinds to one of the looped DNA sites as it did initially. The role of ATP hydrolysis is to prevent the reverse reaction and achieve a unidirectional movement along DNA. The net rate of this cyclic reaction *k*_cycle_ depends on physical quantities, including the rate of ATP hydrolysis by condensin. Although the model proposed by Marko et al. [8] describes the translocation of the bacterial SMC complex along DNA, the basic idea is applicable to chromatin loop formation by other condensins. Hereafter, we refer to these driving force candidates as *motor pulling scenario* and *thermal driving scenario*.

For each scenario, the different energy sources provide different orders of energy to the chromatin fiber for loop formation. As shown in Fig. 1 (b), quantitative estimations of free energy of looped and unlooped chromatin fibers can be used to evaluate the relationship between the energy delivered and the efficiency of loop formation. Thus, the validity of each scenario can be estimated by comparing their efficiencies and by observing the process experimentally in cells.

In this study, we evaluated the free energy difference before and after chromatin loop formation using a polymer chain model, and investigated the source of the driving force for chromatin loop formation and growth. First, we evaluated the free energy associated with loop conformation using a chain model that does not account for excluded volume interaction, which we call the *phantom loop*. Then, we integrated the excluded volume interaction into the free energy difference by employing the mean-field theory (*interacting loop*). By comparing the free energy difference with the energy gain from thermal fluctuation or ATP hydrolysis, we estimated the validity of each scenario.

## Results

### Statistical mechanics of polymer chain model

In this section, we present a face-centered cubic lattice model for polymer chains as a model of the chromatin/DNA fiber. Such fibers differ by their bending stiffness, and the microscopic energy defined by the angle between two consecutive polymer segments regulates this stiffness. In addition, we explain the transfer matrix method used to calculate the free energy of the polymer model without constraints.

### Lattice chain model

A chromatin/DNA fiber is described by a chain model composed of consecutive rod-like segments with the length *b* on a face-centered cubic lattice [9]. Each segment’s direction corresponds to one of the 12 unit vectors of the face-centered cubic lattice; 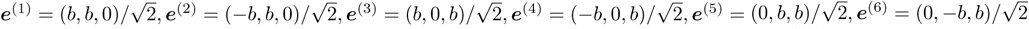 and ***e***^(*η*)^ = −***e***^(*η*−6)^ (*η* = 7, 8, …, 12) (see Figs. 2 (a) and (b)).

**Fig 2.**
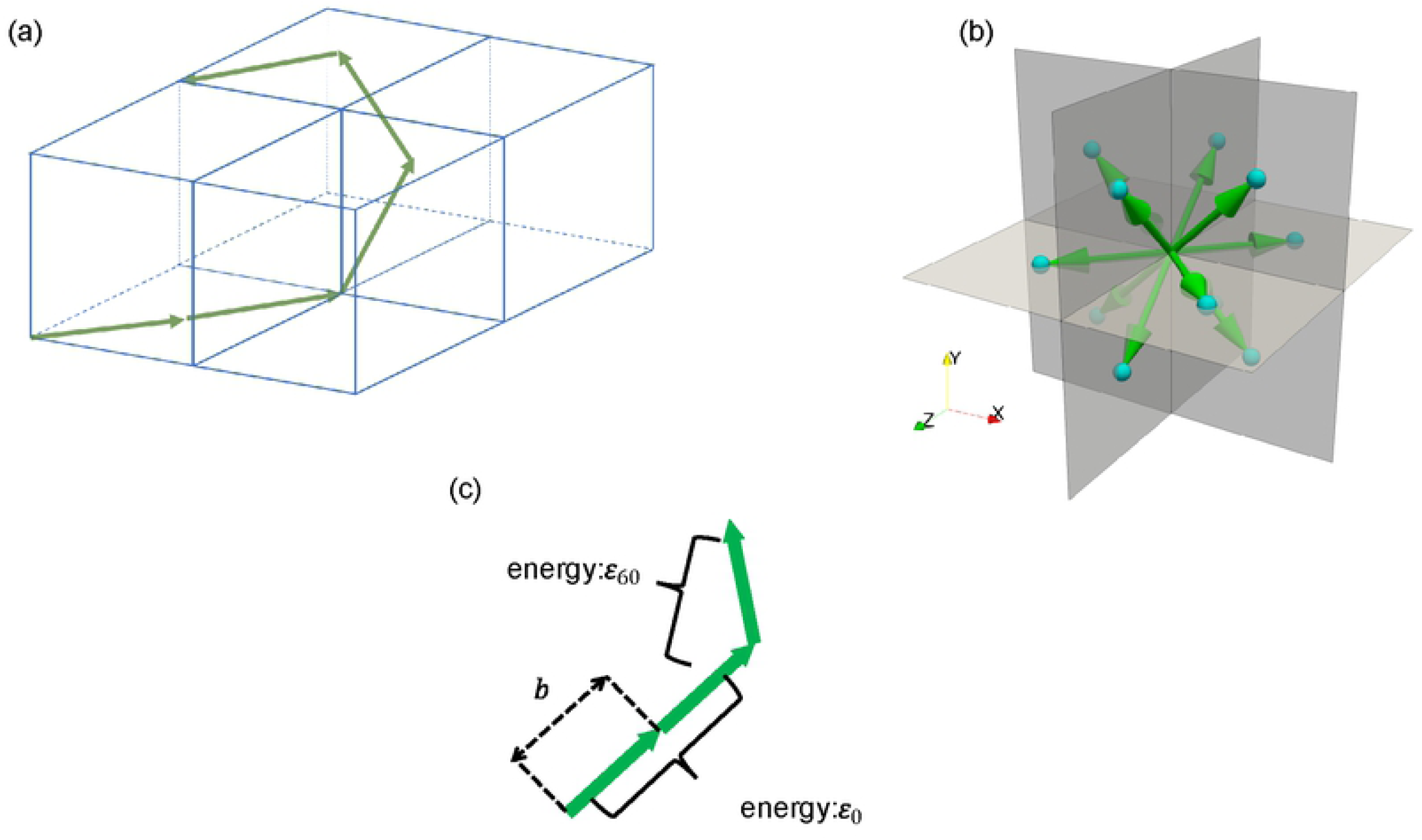
(a) Scheme of the chain model. The model is depicted as walks on the face-centered cubic lattice, where the end of each segment is either at a corner or at a face-center point. (b) Unit vectors of the face-centered cubic lattice. (c) Bending energy of the chain. When the angle between two consecutive segments takes 0° or 60°, the chain obtains the energy *ε*_0_ or *ε*_60_, respectively. *b* is the size of a segment in a chain.

Here, we present the microscopic bending energy for the chain model and we calculate free energy. In our model, two consecutive segments taking 0° and 60° store the microscopic energies *ε*_0_ and *ε*_60_, respectively (see Fig. 2 (c)). Moreover, the storing energy for either of the other angles is *ε*_other_ = ∞, *i.e.* these angles are energetically prohibited. In the scheme of the canonical ensemble in the statistical mechanics, the statistical weights of the angles 0° and 60° are exp [−*ε*_0_/(*k*_B_*T*)] and exp [−*ε*_60_/(*k*_B_*T*)], respectively, where *k*_B_ is the Boltzmann constant and *T* is the temperature. The statistical weight of the other angles becomes exp [−*ε*_other_/(*k*_B_*T*)] = exp [−∞ /(*k*_B_*T*)] = 0. The energy difference Δ*ε* ≡ *ε*_60_ − *ε*_0_ quantitatively describes the chain stiffness. Here, we refer to the relation of two consecutive segments that form an angle of 0° as “parallel state” (or parallel conformation), while the 60° state is termed “bending state” (or bending conformation).

### Transfer matrix method

To calculate the partition function of the chain model with the bending energy, we used the transfer matrix method. We defined 

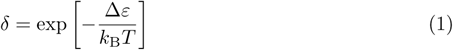

and we introduced the transfer matrix [9] as follows 

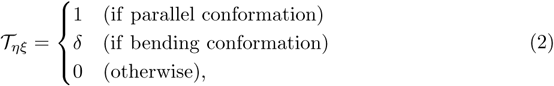

where *η* and *ξ* (*η, ξ* = 1, 2, …, 12) are the indexes of orientation of two consecutive segments. For example, *𝒯*_11_ = 1 represents the statistical weight of the parallel state where both the *n*-th and *n* + 1-th segments point to the direction parallel to ***e***^(1)^. Using eqn. (2), we calculated the partition function of the chain composed of *N* + 1 segments in the free space as follows 

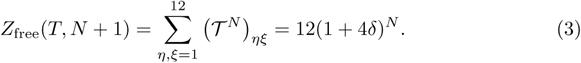

The prefactor 12 means that the orientation of the initial segment can be directed to any of the 12 vectors ***e***^(1)^, ***e***^(2)^ …, ***e***^(12)^. Factor 1 + 4*δ* derives from the fact that two consecutive segments can take one parallel state or one of four bending states.

### Polymer chain model in condensed chromosome

Loop formation restricts the conformation of a polymer chain and thus increases its free energy. Chromosome condensation, along with loop formation, increases the interaction of the excluded volume between loops. This also modifies polymer chain free energy significantly. Here, we included these two modifications on the calculations of polymer chain free energy described above. First, we introduced the loop constraint on the lattice polymer model and developed a calculation method for the free energy difference before and after the loop formation, which is called simply free energy difference. Then, we integrated the effect of the excluded volume interaction using the mean-field theory.

### Loop constraint

We calculated the partition function of a looped polymer chain whose end-to-end distance is equal to zero. Since the conformational free energy calculated by the transfer matrix only refers to segments orientation, we had to integrate the degrees of freedom of each segment ‘s position in order to evaluate the end-to-end distance. As the end-to-end vector of the chain is obtained by the sum of the *N* + 1 segment vectors, we calculated the statistical weight 

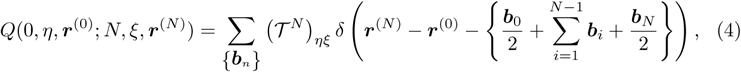

where the 0-th segment at the position ***r***^(0)^ and the *N* -th segment at the position ***r***^(*N*)^ point to ***e***^(*η*)^ and ***e***^(*ξ*)^, respectively. By using the Fourier expression of *δ* function, eqn. (4) is rewritten as 

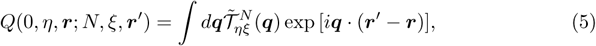

where, 

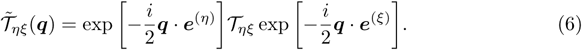

The loop structure of the chain is obtained from the chain conformation under a constrained condition where each end of the chain has the same position. Therefore, the partition function of the loop structure of a chain composed of *N* + 1 segments, *Z*_loop_(*T, N* + 1), is calculated as 

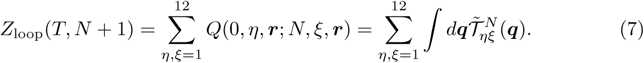

The free energy difference of a phantom loop, Δ*F*_0_, is calculated as 

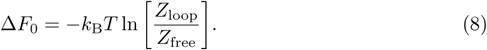

The free energy difference of multiple phantom loops is also calculated as 

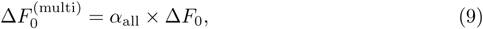

where each loop is assumed to be composed of *N* + 1 segments and *α*_all_ is the number of loops in the system.

### Excluded volume interaction

We integrated the excluded volume interaction into the free energy of the polymer model. We adopted the mean-field theory, in which the statistical properties of multiple interacting loops were determined using approximate calculations of a single loop under a potential field. In general, to introduce the excluded volume interaction between segments using the mean-field theory, we start with a Hamiltonian 

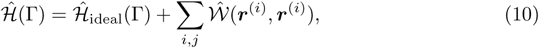

where the first term describes the Hamiltonian of a single polymer without excluded volume interaction (an ideal system) and the other term refers to the interaction between segments. Moreover, 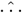 means that the quantity · · · is a function of phase space Γ = {***r***^(0)^, ***b***^(0)^, ***b***^(1)^, · · ·, ***b***^(*N*)^}. The expression (10) is rewritten as 

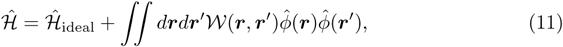

where *ϕ*(***r***) is the segment density field defined as 

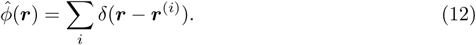

We assume that the spatial correlation in the interaction term of Hamiltonian (11) is negligible, 

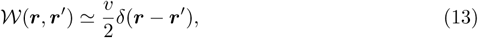

where *v* is the excluded volume parameter and the prefactor 1/2 avoids double-counting of the interaction. Then, Hamiltonian (11) becomes 

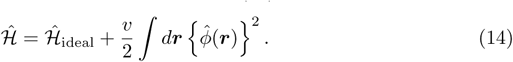

The partition function of this Hamiltonian is 

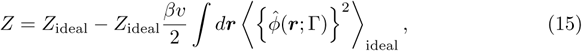

where *βv* is regarded as the perturbation parameter. ⟨… ⟩_ideal_ means the ensemble average value of … over the ensembles of the ideal system and *Z*_ideal_ is the partition function of the ideal system. Free energy is derived from eqn. (15) as follows: 

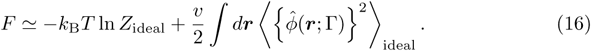

Here, for simplicity, we assumed that the segment density is spatially uniform: 

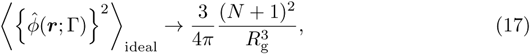

Here, *R*_g_ is the gyration radius of the polymer chain, defined as 

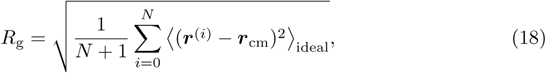

where ***r***_cm_ is the center of mass of the chain: 

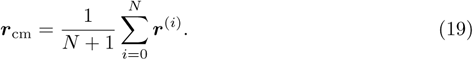

Note that in the mean-field theory, the gyration radius in (17) is averaged over the ensemble of the ideal system.

By replacing each of the *Z*_loop_ and *Z*_free_ by *Z*_ideal_ in eqn. (16), we obtained the free energies of the interacting loop and the chain model in the free space, respectively. Then, we calculated the free energy difference Δ*F* as follows. 

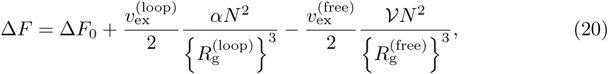

Where 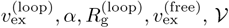 and 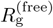 are the excluded volume parameter of a single loop, the number of loops interacting with each other, the gyration radius of the phantom loop, the excluded volume parameter of the model chain in the free space, the volume of the system, and the gyration radius of the model chain in the free space, respectively. It should be noted that the integration interval of the interaction term in eqn. (16) gives us the volume 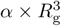. For simplicity, we have set 

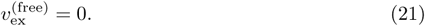

Thus, we obtained the free energy difference of the multi-loop system, 

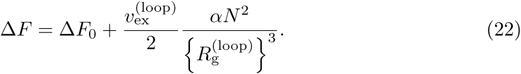

In the discussion, for the sake of simplicity, 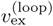 is denoted by *v*_ex_.

### Phantom loop structure

Here, the chromatin/DNA fiber is characterized by the chain stiffness and stiffness is described by the persistence length *l*_p_ as follows 

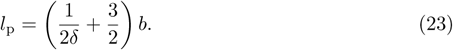

This expression is obtained by fitting the segment orientation correlation along the chain. The details of the fitting are shown in *Material and Method*. We selected a persistence length of chromatin of 30nm [10]. Following [3], we set the segment size to *b* = 10 nm, which specifies the minimum spatial scale of the system. Additionally, bare DNA can also be described by our model if we select *l*_p_ = 50nm, which is the value of DNA [11].

We calculated the free energy difference of a single phantom loop, which corresponds to a chromatin loop. As shown by the purple line in Fig. 3, Δ*F*_0_ is the increasing function of *N* and the slope decreases with *N*. In fact, the free energy difference between the two chromatin states is well fitted by

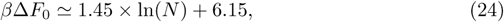

where the non-linear least squares method is used for fitting.

We estimated the efficiencies of the two driving force scenarios for loop growth, under the phantom loop model. One of the scenarios is that of thermal driving, where the source of the driving force is thermal fluctuation. Under thermal fluctuation, a physical model typically gains on an average the energy 1*k*_B_*T* from thermal noise. Here, we supposed that a chromatin loop with contour length *N* is held by condensin and its length is forced to increase owing to thermal fluctuation. The typical increase in length 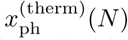 is calculated based on the thermal energy gained 1*k*_B_*T* (see the two purple dashed lines in Fig. 3). 

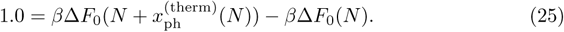

**Fig 3.**
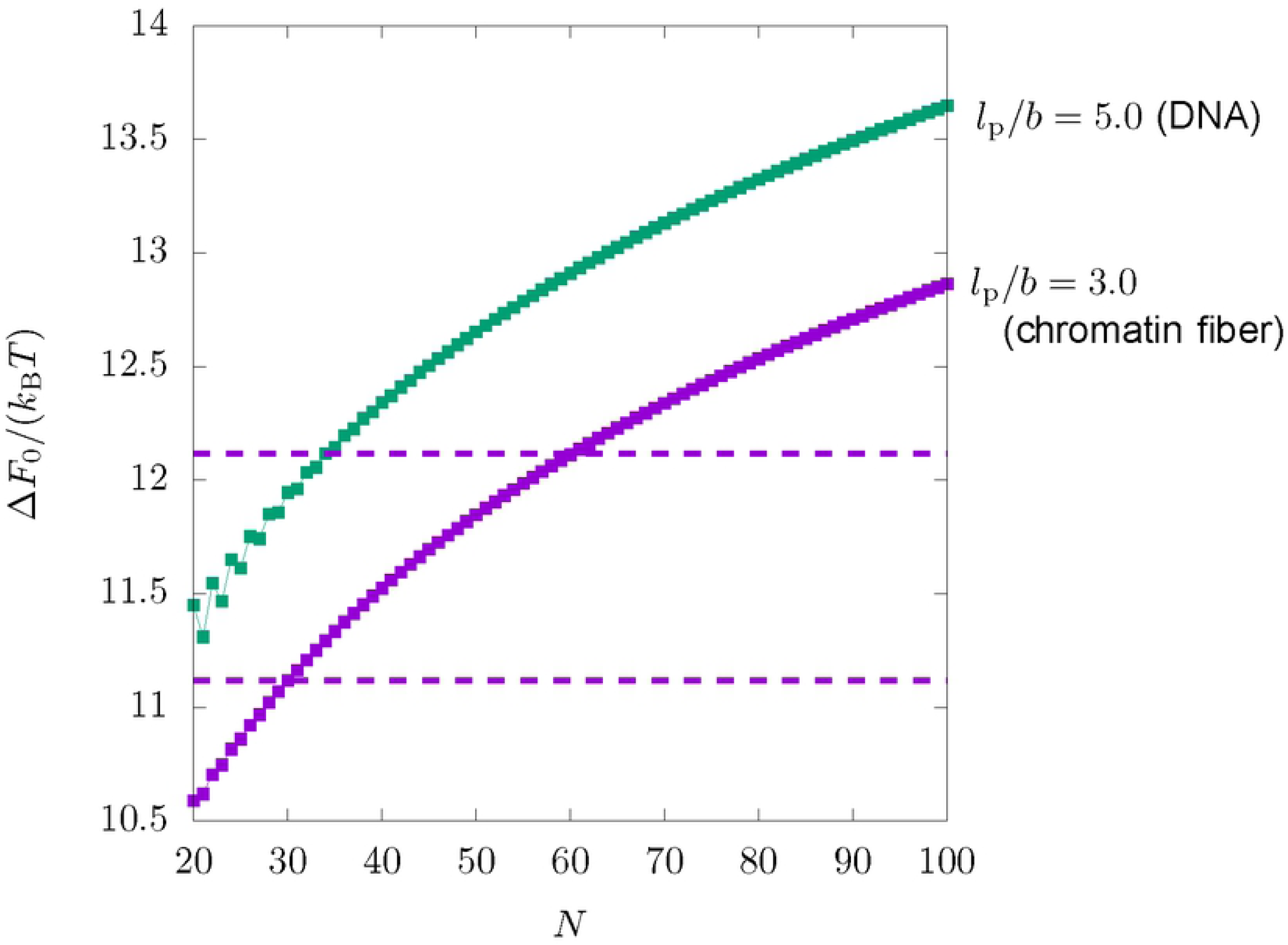
Free energy difference of a phantom loop. The vertical and horizontal axes represent the non-dimensional free energy difference Δ*F*_0_/(*k*_B_*T*) and the number of segments in the loop *N*, respectively. The purple and green symbols represent the free energy difference of the chromatin fiber and that of bare DNA. The difference between these polymers is identified by the value of the persistence length *l*_p_. The two purple dashed lines represent the values of Δ*F*_0_(*N* = 30, *l*_p_*/b* = 3.0)/(*k*_B_*T*) and (Δ*F*_0_(*N* = 30, *l*_p_*/b* = 3.0) + *k*_B_*T*)/(*k*_B_*T*).

Note that the contribution of the constant to Δ*F*_0_ disappears in eqn. (25). Using eqn. (23), we obtained 

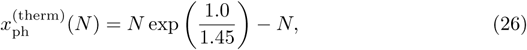

which is the loop-growth efficiency of the thermal driving scenario. In the case of *N* = 30 the typical increase in length is 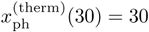. In the motor pulling scenario, loop growth is coupled with ATP hydrolysis, which releases about 12.2*k*_B_*T*. Experiments in the chicken cell line DT40 showed that the typical contour length of loops in matured chromosome is about 24 *µ*m and that the duration of loop formation (from the beginning of prophase until the end of prometaphase) is about 60 min (= 3600 s). [5]. The ATP hydrolysis rate of condensin in a *Xenopus* cell is *k*_ATP_ = 0.90 [12]. Thus, the number of ATP molecules used during loop formation is 

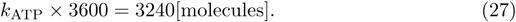

Using eqn. (23) and the number of ATP molecules (27), we can obtain the loop-growing efficiency of the motor pulling scenario as follows 

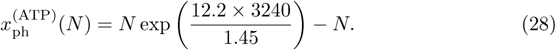

When *N* = 30, we obtain 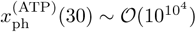. As the segment size is 10 nm, the contour length of the loop can grow from 300 nm to 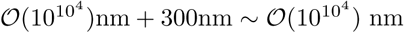.

Considering the estimations above, one can easily notice that the motor pulling scenario is not suitable for describing the real loops that form in chromosomes. Based on such estimations and the mitotic conditions known in DT40 cells, we examined the validity of the loop-growth efficiency of the thermal driving scenario. According to this scenario, ATP hydrolysis should occur within one reaction cycle. Therefore, the typical time scale of one cycle can be approximated as 

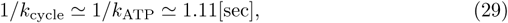

where we used the ATP hydrolysis rate of condensin measured in a *Xenopus* cell *i.e. k*_ATP_ = 0.90 [12]. Thus, the number of cycles during loop formation is given by 

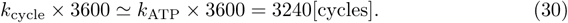

To achieve a loop with a contour length of 24.0 × 10^3^[nm], the average increase in length during loop growth must be 

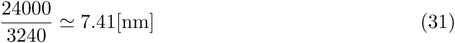

which is less than the estimated loop-growth efficiency 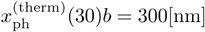. In both scenarios, the loop-growth efficiency estimated by our model is too high to fit the actual loop growth that occurs in cells. This indicates that the phantom loop model is not suitable to describe the formation of loops in chromosomes. As the free energy difference of bare DNA is described as a parallel displacement of that of chromatin (see Fig. 3), the considerations made above also apply to DNA.

### Interacting loop structure

We showed the free energy difference of the multi-loop system including the excluded volume interaction by employing the mean-field theory (interacting loop model). In this model, the free energy difference Δ*F* depends not only on *N* and *l*_p_ but also on the parameters *α* and *v*_ex_, in contrast with the phantom loop model.

To obtain the essential behavior of the free energy difference, we first calculated free energy difference considering fixed *α* = 20 and *βv*_ex_ = 5.00 × 10^−2^. *α* is estimated by the number of loops overlapping each other according to the chromosome conformation reported by Gibcus *et al.* [5] (see *Materials and Methods* for more details). As shown in Fig. 4 (purple line) the free energy difference of the interacting chromatin loop increases linearly with the number of segments within a large *N* range(*N >* 30).

**Fig 4.**
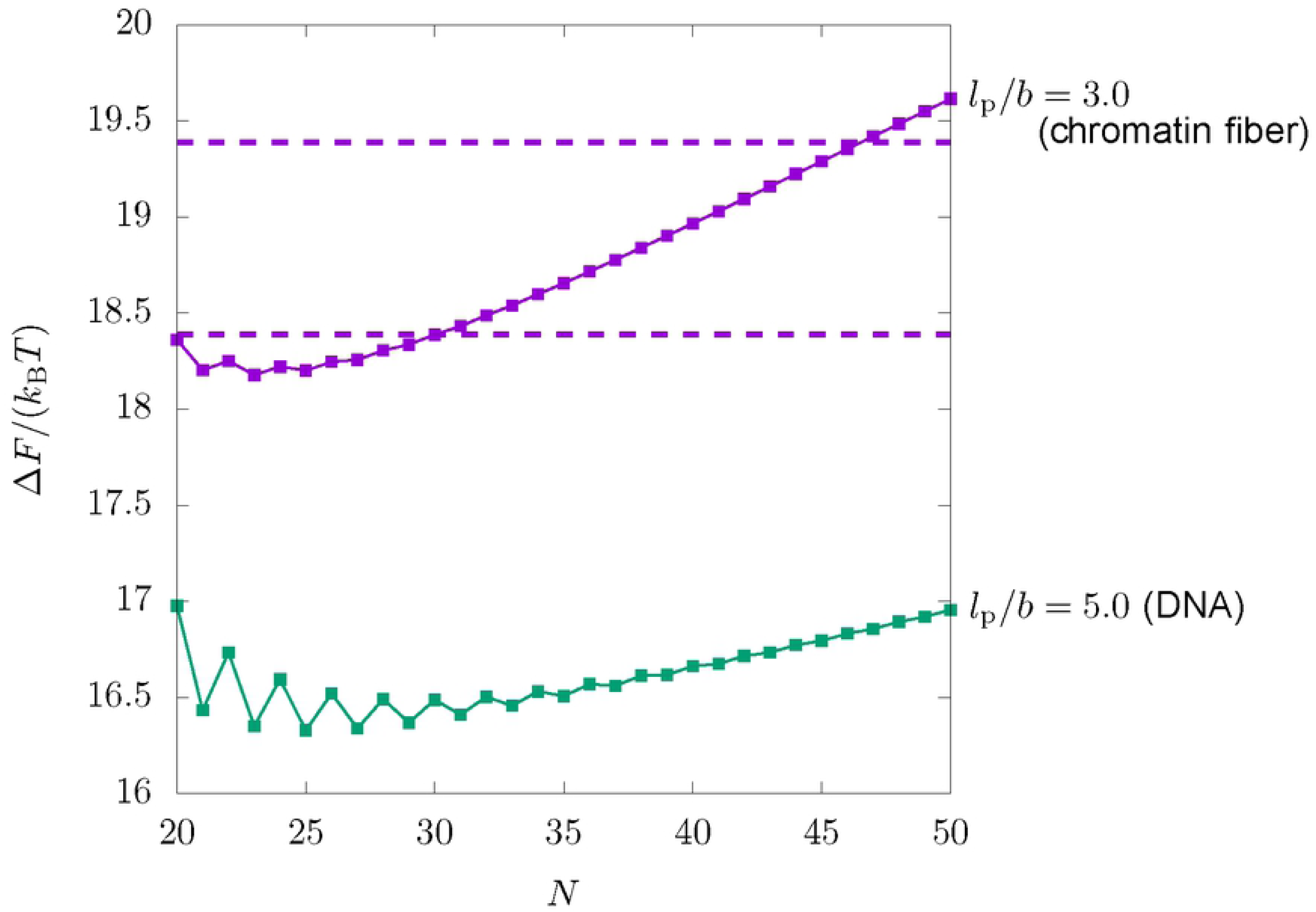
Free energy difference of interacting loop considering *α* = 20 and *βv*_ex_ = 5.00 × 10^−2^. The vertical and horizontal axes represent the free energy difference Δ*F/*(*k*_B_*T*) and the number of segments in the loop *N*, respectively. The purple and green symbols depict the free energy differences of the interacting chromatin loop and of the interacting bare DNA loop, respectively. The two purple dashed lines represent the free energy differences when *N* = 30 and when 1 *k*_B_*T* is added to this value.

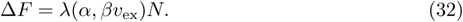

We determined the linear coefficient as follows 

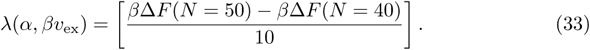

In the case of *α* = 20 and *βv*_ex_ = 5.00 × 10^−2^, *λ*(*α, βv*_ex_) ≃ 15.

By using the free energy difference of the interacting loop model, we evaluated the validity of the thermal driving and the motor pulling scenario. As the free energy difference behaves as a linear function of *N*, the increase in segments of the chromatin interacting loop using thermal fluctuation, 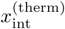, was estimated from the slope of the free energy difference plot. Here, 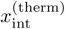 is defined as 

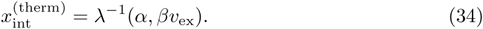

The typical increase in length depends on the excluded volume parameter *βv*_ex_, which is shown in Fig. 5.

**Fig 5.**
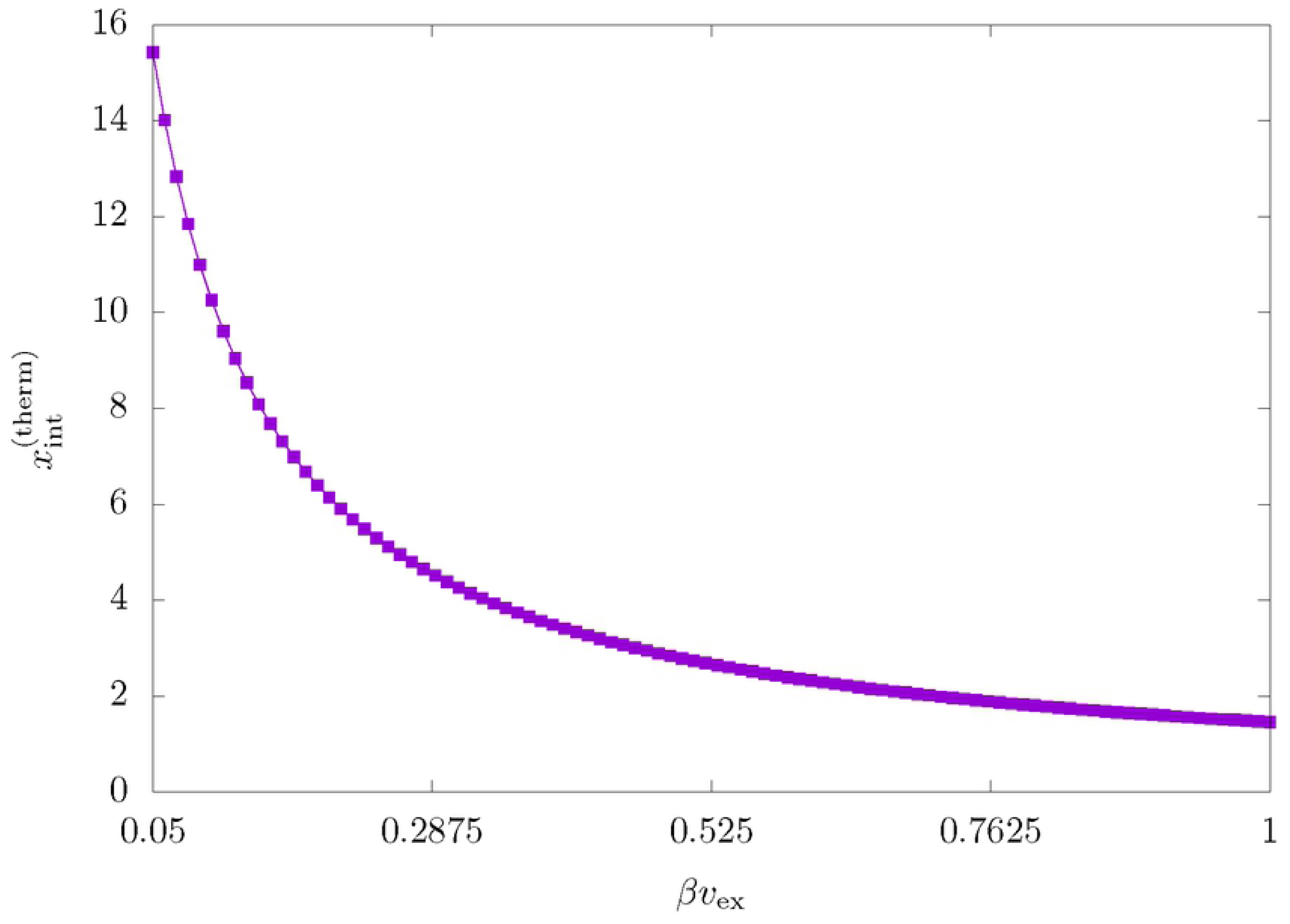
Behavior of 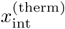 with *βv*_ex_. The number of loops *α* = 20 is fixed.

As the energy gained from the ATP hydrolysis is about 12.2*k*_B_*T* and the number of ATP molecules is 3,240, 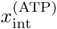 was estimated as 

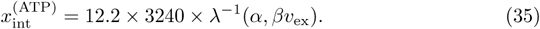

The behavior of 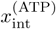 with *βv*_ex_ is shown in Fig.6.

**Fig 6.**
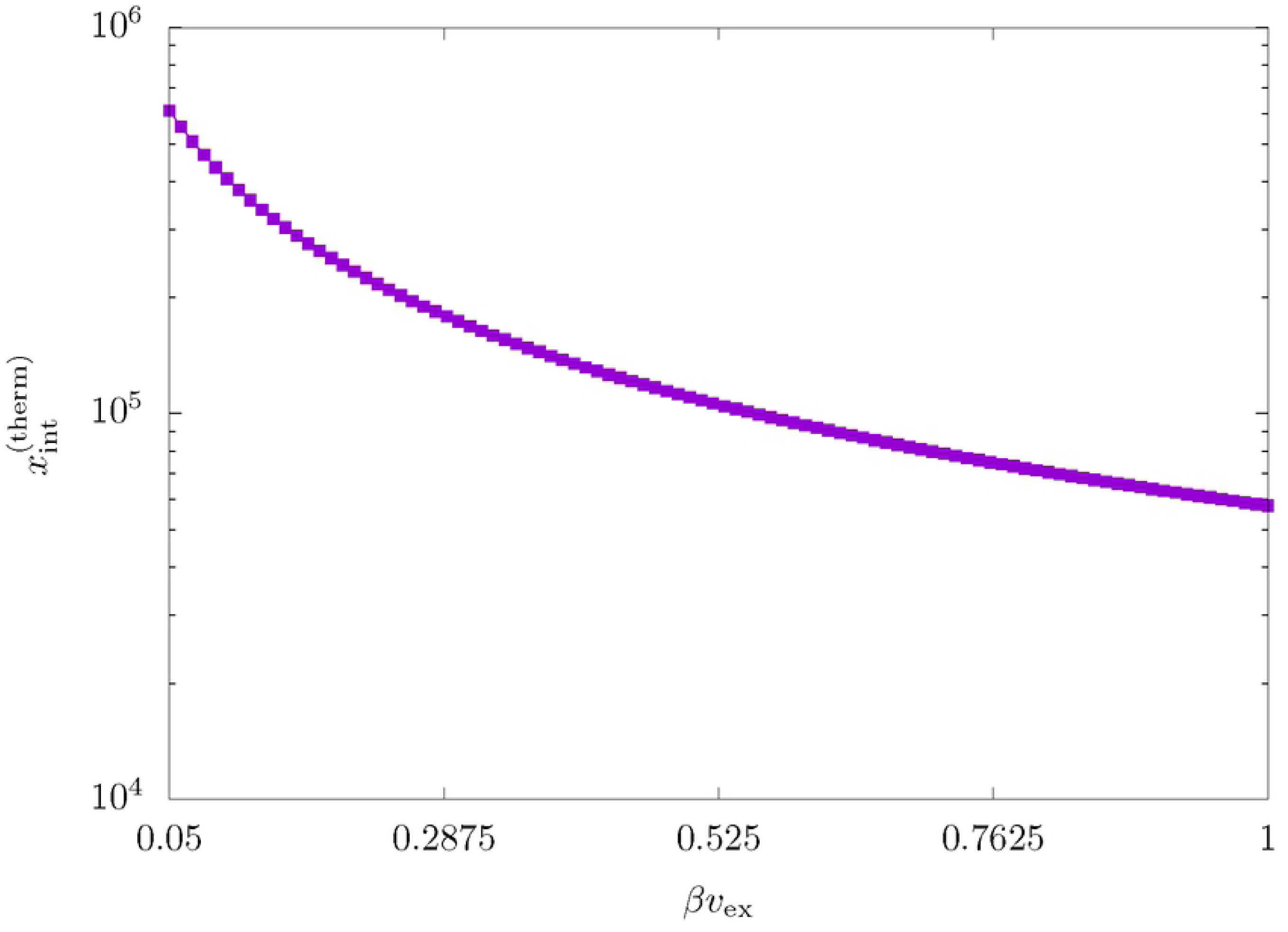
Behavior of the 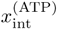 of chromatin with *βv*_ex_ when the number of loops *α* = 20 is fixed.

Based on these results (Figs. 5 and 6), we evaluated the excluded volume parameters *βv*_ex_ for both scenarios. For the thermal driving scenario, as shown in eqn. (31), the loop is estimated to grow 7.47 [nm] (corresponding to 0.747 × *b*) during each cycle. By applying such estimation to the interacting loop model (Fig. 5), the excluded volume parameter is expected to be higher than one 

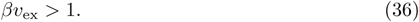

According to the motor pulling scenario, the number of segments in the matured chromatin loop is 2,400 (24 × 10^3^ [nm]). Therefore, the excluded volume parameter (Fig. 6) is expected to be much higher than one 

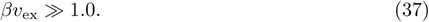

Note that for both scenarios, the excluded volume parameters are estimated to exceed one. This represents a break of the perturbation theory as this is an expansion parameter. Such possibility will be addressed further in the discussion.

## Discussion

In our study, we employed the lattice polymer model to describe a chromatin fiber and determine the free energy difference between its unlooped and looped state. Moreover, the effect of the excluded volume was included using the mean-field theory. Using this strategy we evaluated the validity of two possible scenarios for the driving force of loop formation: motor pulling scenario and thermal driving scenario. First, we confirmed that the excluded volume interaction is an essential resistance effect during loop formation in both scenarios. Our estimations suggest that if thermal fluctuation is the main driving force of loop growth, the dimensionless excluded volume parameter *βv*_ex_ should be higher than one, *i.e., βv*_ex_ *>* 1. Likewise, we propose that if the energy gained from ATP hydrolysis is the source of this driving force, then the excluded volume should be included in *βv*_ex_ ≫ 1.0.

It is worth clarifying the physical meaning of the excluded volume interaction parameter *v*_ex_. In the context of polymer physics, the value of *v*_ex_ is determined by the interaction between segments [13]. *βv*_ex_ *>* 0 describes the repulsive force between segments within the chromatin loop that leads to the wide spatial distribution of segments. This repulsive force is a consequence of the intraloop repulsive interaction overcoming the interloop repulsive interaction. On the other hand, *βv*_ex_ *<* 0 represents the attractive force between segments within the chromatin loop, in which the spatial distances between segments shrinks. This attractive force comes from the interloop repulsion overcoming the intraloop repulsion. *βv*_ex_ = 0 corresponds to a situation where interloop and intraloop repulsions balance out. As shown in *Result*, the situation where *βv*_ex_ *>* 0 is more suitable than the phantom loop case (*βv*_ex_ = 0). This result implies a spatially wide distribution of segments in the chromatin loop due to the repulsive forces associated with loop growth.

Although our perturbation theory on the excluded volume interaction takes a two-body correlation into account, a multi-body correlation (higher than two) can be incorporated to provide an essential effect on this interaction. Supposing that most interactions between segments are repulsive, assuming a multi-body correlation results in a wider spatial distribution of segments and increases the resistance effect against loop growth. By incorporating the multi-body correlation the loop-growth efficiencies, 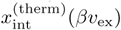 and 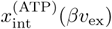, would become more accurate and lower than those previously determined (Figs. 5 and 6). The difference between the calculated and the most accurate loop-growth efficiencies should be slight in the region *βv*_ex_ ≪ 1.0 and considerably large in the region *βv*_ex_ ≥ 1.0. Since *βv*_ex_ is the expansion parameter, the large difference in the latter region indicates the breaking of the perturbation theory. Therefore, we used the perturbation theory to estimate *βv*_ex_, which leads to the breaking of the perturbation. However, the previous remarks about the effect of the multi-body correlation can lead us to another valid conjecture. If in the thermal driving scenario the accurate loop-growth efficiency is similar to the value determined for the region *βv*_ex_ *<* 1.0 and lower than that of region *βv*_ex_ ≥ 1.0, the value of the excluded volume interaction parameter satisfying 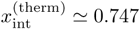 should be 

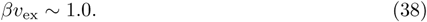

On the other hand, under the motor pulling scenario, the accurate loop-growth efficiency satisfying 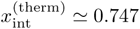 should still indicate *βv*_ex_ ≫ 1.0. Therefore, based on our calculations, we conclude that the value of *βv*_ex_ determines which of the two scenarios is more plausible.

Let us confirm the existence of effective multi-body correlation in vivo condition referring [5]. In this conformation, a 12-Mbp nucleotide sequence has an helical conformation with a contour length of 250 nm and a radius of 100 nm (see *Material and Method*). The nucleotide density in the chromosome is 12 × 10^6^/(*π* × 100^2^ × 250) ∼ 1.6[bp/nm^3^]. This corresponds to 1,600 bp in a cubic in which each edge is 10 nm (segment size). As the size of two nucleosomes is 300 bp, which corresponds to 10nm, 1,600 bp in a cubic correspond to five segments in the occupied volume of a single segment. Therefore, in this case, the effective multi-body correlation is expected. The interaction between nucleosomes affects the environment surrounding the chromatin fiber during loop growth. Such an environment determines the extent and dynamics of loop growth. Therefore, it would be worth to measure the excluded volume interaction between nucleosomes.

Our estimations implicitly assume that the loop grows during all reaction cycles. However, particularly in the thermal driving scenario, the loop does not necessarily need to enlarge at all cycles and such growth can be probabilistic. In this scenario, the two nucleosomes are bound by condensin due to thermal fluctuation; then, one of the nucleosomes is kept bound and the other is released. Although this nucleosome dynamics is coupled with a cycle of chemical reactions including ATP hydrolysis, such chemical reactions can occur in the absence of chromatin capture or nucleosome release by condensin. In such a situation, the estimation of 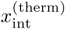 should be modified. For example, if the probability of a loop growth event is 0.0250, the contour length associated with one such event (not during one cycle) should be 7.47/0.025[nm]≃300[nm], which is consistent with the loop-growth efficiency predicted by our phantom loop model. Unfortunately, loop growth probability cannot be quantified because it depends on many physical parameters such as the chromatin binding affinity of condensin and the mobility of chromatin within the cell. To verify in detail the thermal driving scenario, it is important to know the values of these parameters.

Regarding the motor pulling scenario, we implicitly assumed that a single condensin complex is present on the chromatin fiber during looping. However, more than one condensin can be involved in loop formation [3, 14]. In fact, the numbers of condensin I and condensin II can reach the hundreds and the tens of thousands, respectively [14]. In this case, the motor activity of each condensin may cooperate or compete with each other. According to Figure. 6 there are two hypotheses for the cooperation/competition of condensins’ motor activities. In the first hypothesis, the condensins may cooperatively work, in which case the loop-growth efficiency increases. Therefore, inequality (37) is not modified. In the other hypothesis, condensins compete with each other and consequently the loop-growth efficiency decreases. This means that inequality (37) should be modified, for example *βv*_ex_ ≫ 1 → *βv*_ex_ *>* 1. As to the precise value of the excluded volume parameter, the key factor to determine it is the cooperative activity of condensins.

One limitation of our study is that we do not discuss the free energy difference when the chain length *N* is small. Because the lattice model assumes a large number of segments, our model does not apply when the *N* is small. For example, the free energy difference of bare DNA when considering a small *N* reflects the discrete characteristics of the lattice polymer, whose features do not reproduce the physical properties of DNA (Fig. 4, green line). Thus, the earliest steps of loop formation fall outside the range of our model. To study the early behavior of chromatin, we need to create a different model that calculates the free energy difference between unbent and bent rods based on mechanics rather than on statistical mechanics. Studying the loss of free energy associated with a small *N* may be important to understand chromosome condensation because such loss occurs close to the region where condensin binds. Therefore, free energy loss may contribute to chromatin dynamics during the early stages of chromosome assembly. A model applicable to a small *N*, that is able to describe these initial stages, should be focus of future work.

## Materials and methods

### Path integral of model chain

We derived the path integral expression eqn. (5) as follows: 

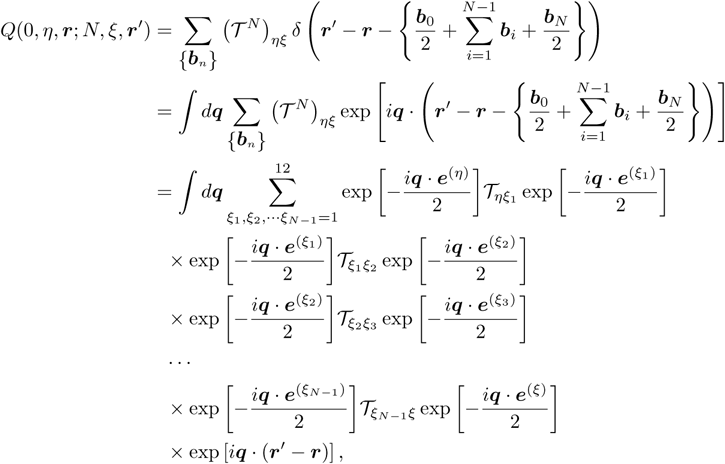

where we took into account that the *i*-th segment vector ***b***_*i*_ can be any of the 12 basic lattice vectors. Therefore, 

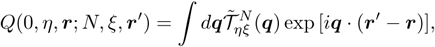

where, 

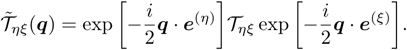

### Perturbation theory and mean field theory

Here is shown in detail how we derived eqn. (20). The expression (10) was rewritten as 

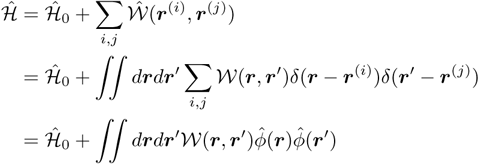

We assumed that the spatial correlation in the interaction term of the Hamiltonian (11) was negligible, *i.e.* 

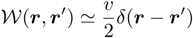

Then, the Hamiltonian (11) was rewritten as eqn. (14). The partition function was obtained from this Hamiltonian (14) as 

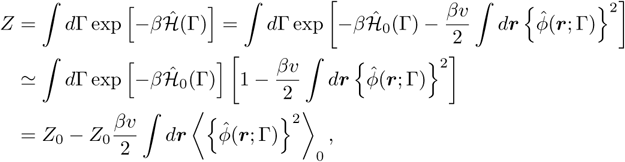

where *βv* is regarded as the perturbation parameter. The free energy is derived as follows: 

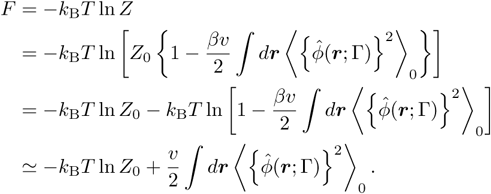

### Determining of expression of persistence length

To determine the expression the orientation correlation along the chain was computed as the persistence length is defined by the contour length of the polymer chain when the orientation correlation of the chain is lost. The orientation correlation function *G*_ori_(*j*) was described as 

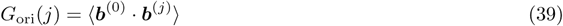

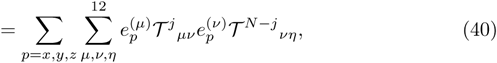

where *j* is the segment index and *p* denotes Cartesian coordinates *x, y* or *z*. The persistence length satisfies 

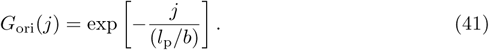

By using the transfer matrix, *G*_ori_(*j*) in the case of *δ*^−1^ = 3.00 and *δ*^−1^ = 8.00 are calculated as shown in Figs. 7 and 8

**Fig 7.**
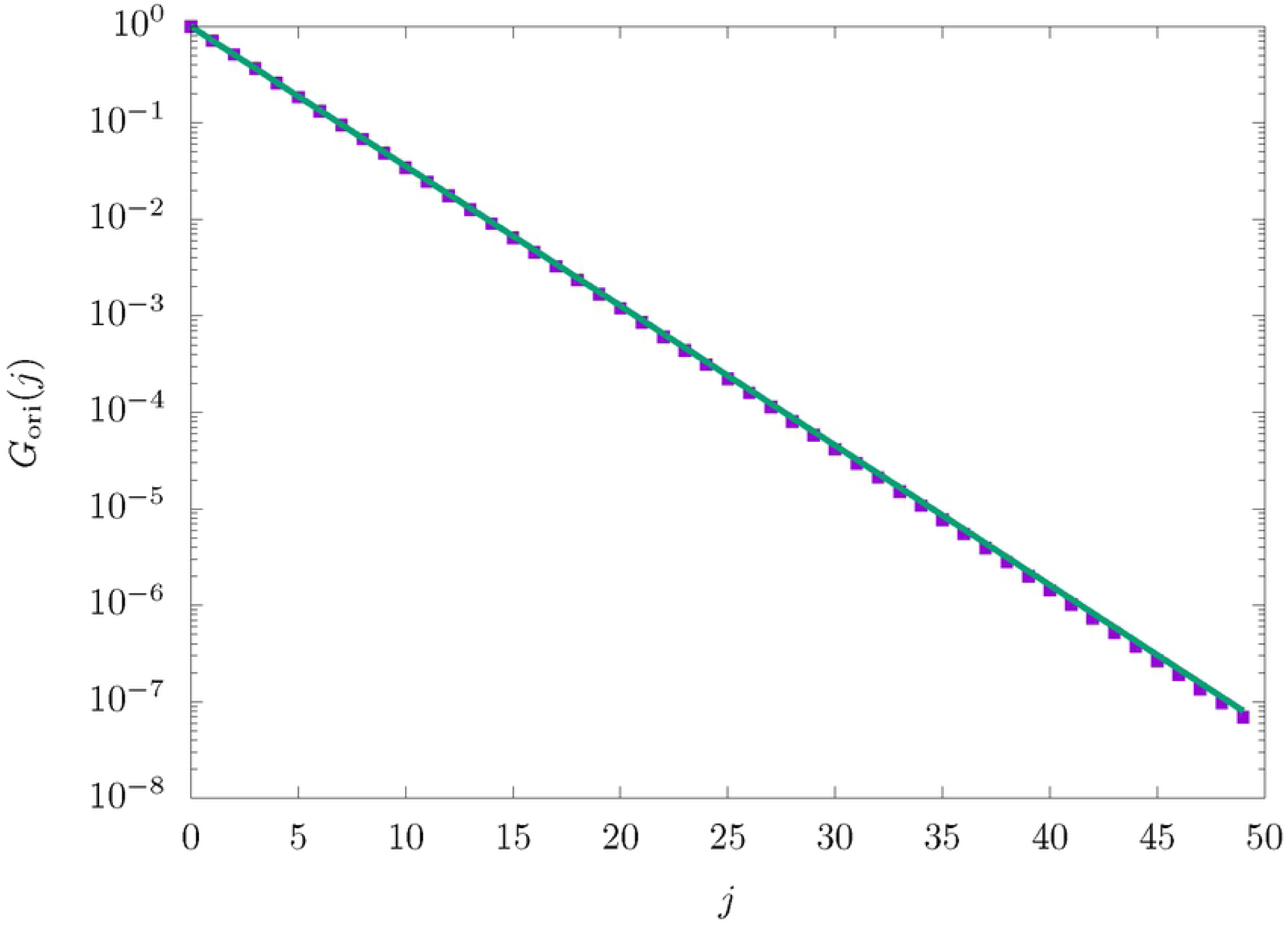
Behavior of *G*_ori_(*j*) in the case of *δ*^−1^ = 3.00 (corresponding to*l*_p_*/b* = 3.00). The purple symbol and the green line represent the numerical result and the fitting function eqn. (42), respectively.

**Fig 8.**
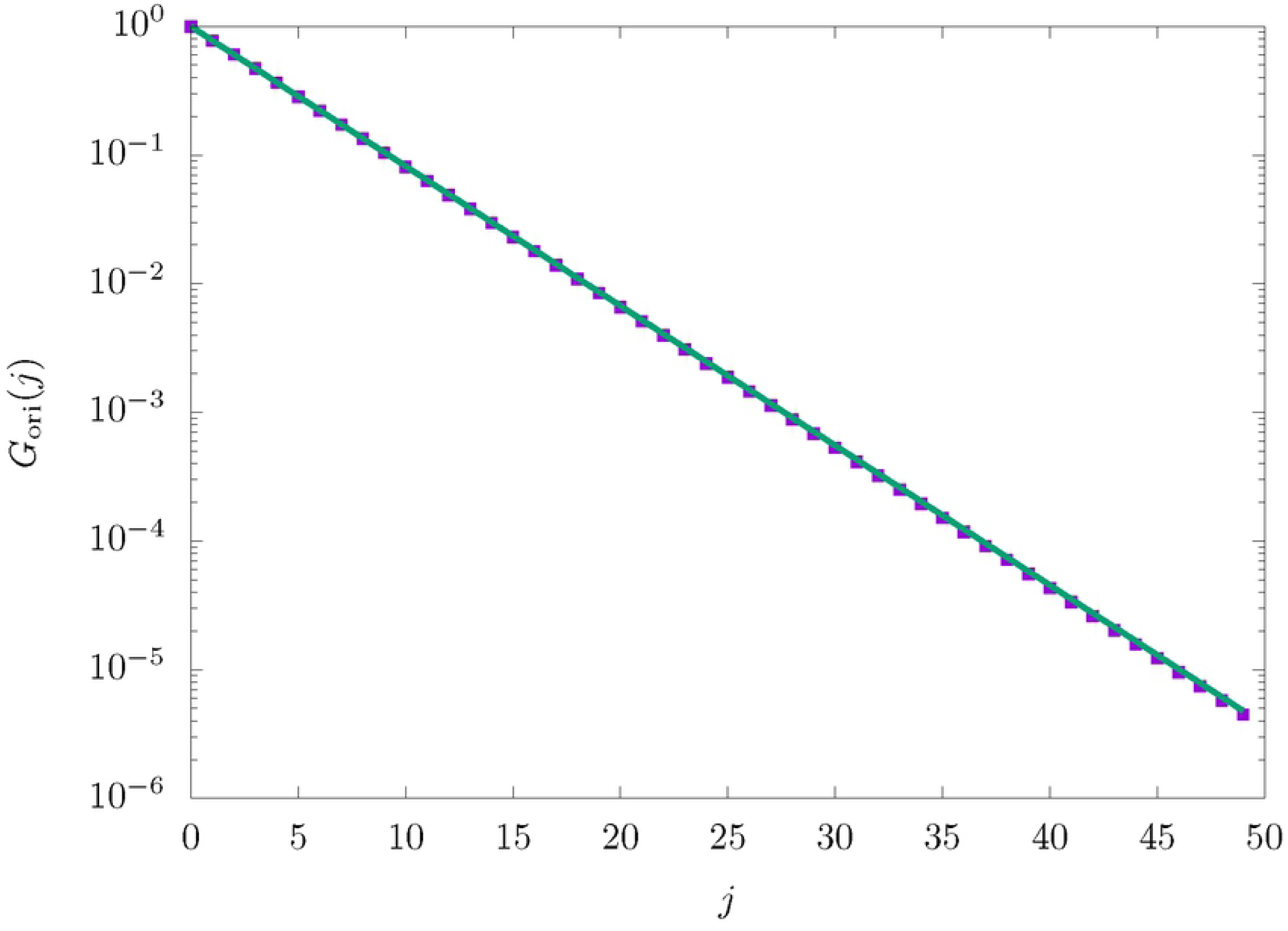
Behavior of *G*_ori_(*j*) in the case of *δ*^−1^ = 8.00 (corresponding to *l*_p_*/b* = 5.00). The purple symbol and the green line represent the same as in Fig. 7.

These numerical results are fitted by

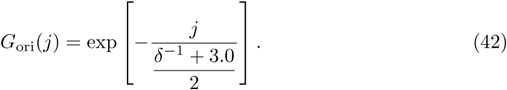

By using eqns. (41) and (42), the persistence length *l*_p_ was described as 

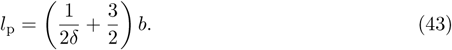

When the value of *b* was not strictly determined, the constant contribution to *l*_p_ is renormalized in the segment size *b* (for example, in the case where the value of *b* simply specifies the order of the size of the monomer in the polymer [9]). Therefore, in such a case, the persistence length can be defined as *l*_p_ = *b/δ*. However, in this study we cannot do so because the spatial scale is clearly determined by the value of *b, i.e., b* = 10[nm]. Here, we have to use eqn. (43) to define the persistence length.

### Gyration radius

The square of the gyration radius 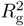 was defined as

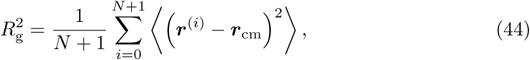

where ***r***_g_ is the center of mass: 

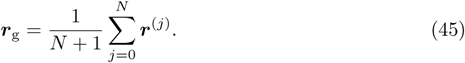

Equation (44) was rewritten as

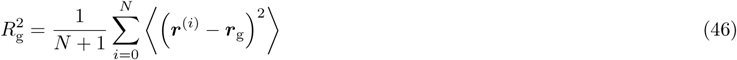

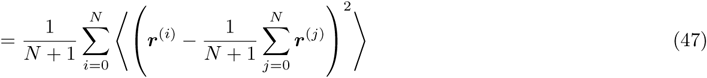

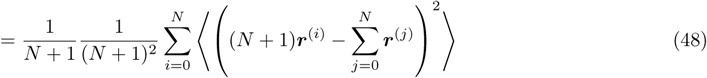

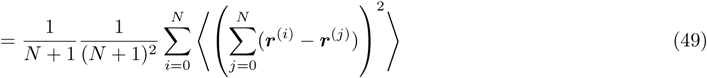

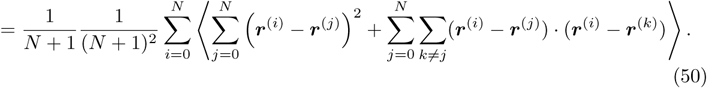

### Estimation of *α*

The number of loops interacting with each other *α* were estimated based on the chromatin conformation shown in Fig. 9. We denoted the distance between the centers of mass of the nearest neighbor loops by Δ. As the gyration radius of the loop is *R*_g_,

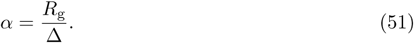

**Fig 9.**
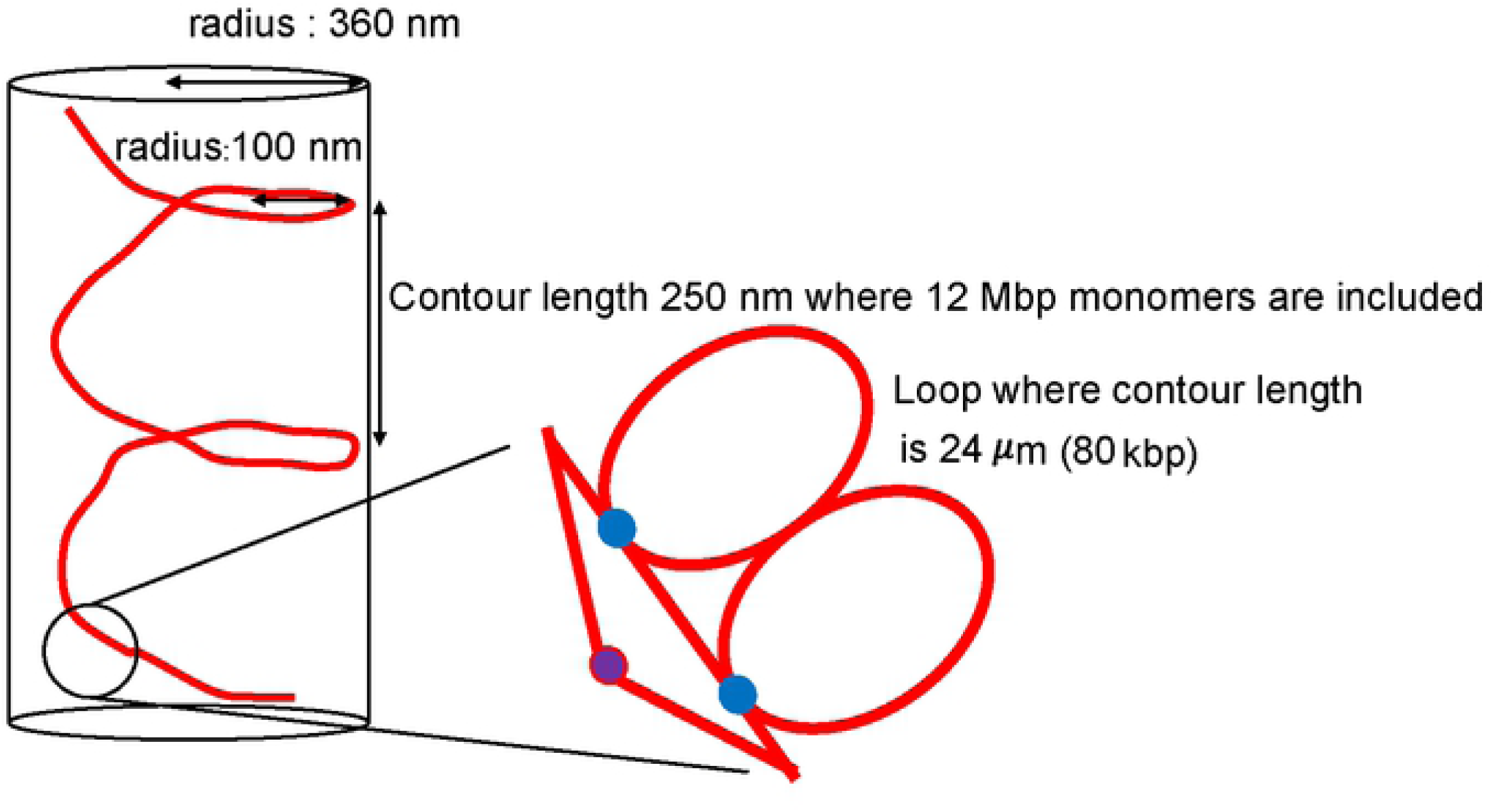
Chromosome structure reported by Gibcus *et al.* [5], based on Hi-C data and simulation studies. The red line represents a chromosome with a helical conformation including a nested loop structure. The blue and purple circles depict condensin I and II, respectively.

We calculates the number of loops including the spatial range Δ. During prometaphase, the contour length of each loop is 80kbp. Additionally, 1 bp=0.3 nm which leads to

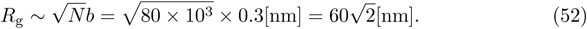

As 150 loops exist in one pitch of the helical conformation shown in Fig. 9,

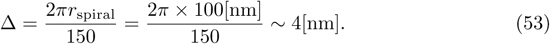

The number of interacting loops *α* is calculated as

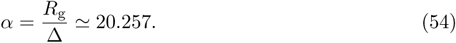

Therefore, we fixed *α* = 20.

## Acknowledgments

This work was supported by a Grant-in-Aid for Scientific Research (grant number 18H05276) from The Ministry of Education, Culture, Sports, Science and Technology (MEXT), Japan. Numerical computations were partially performed using RIKEN’s supercomputer system HOKUSAI GreatWave.

**Supporting information**

## References

1. Sakai Y, Mochizuki A, Kinoshita K, Hirano T, Tachikawa M. Modeling the functions of condensin in chromosome shaping and segregation. PLoS computational biology. 2018;14(6):e1006152.

2. Alipour E, Marko JF. Self-organization of domain structures by DNA-loop-extruding enzymes. Nucleic acids research. 2012;40(22):11202–11212.

3. Goloborodko A, Imakaev MV, Marko JF, Leonid M. Compaction and segregation of sister chromatids via active loop extrusion. eLife. 2016;5.

4. Terakawa T, Bisht S, Eeftens JM, Dekker C, Haering CH, Greene EC. The condensin complex is a mechanochemical motor that translocates along DNA. Science. 2017;358(6363):672–676. doi: 10.1126/science.aan6516.

5. Gibcus JH, Samejima K, Goloborodko A, Samejima I, Naumova N, Nuebler J, et al. A pathway for mitotic chromosome formation. Science. 2018;359(6376). doi: 10.1126/science.aao6135.

6. Ganji M, Shaltiel IA, Bisht S, Kim E, Kalichava A, Haering CH, et al. Real-time imaging of DNA loop extrusion by condensin. Science. 2018;360(6384):102–105. doi: 10.1126/science.aar7831.

7. Goloborodko A, Marko JF, Mirny LA. Chromosome Compaction by Active Loop Extrusion. Biophysical Journal. 2016;110(10):2162–2168. doi: https://doi.org/10.1016/j.bpj.2016.02.041.

8. Marko JF, De Los Rios P, Barducci A, Gruber S. DNA-segment-capture model for loop extrusion by structural maintenance of chromosome (SMC) protein complexes. Nucleic Acids Research. 2019;47(13):6956–6972. doi: 10.1093/nar/gkz497.

9. Yokota H, Kawakatsu T. Modeling induction period of polymer crystallization. Polymer. 2017;129:189–200. doi: https://doi.org/10.1016/j.polymer.2017.09.022.

10. Cui Y, Bustamante C. Pulling a single chromatin fiber reveals the forces that maintain its higher-order structure. Proceedings of the National Academy of Sciences. 2000;97(1):127–132. doi: 10.1073/pnas.97.1.127.

11. Lu Y, Weers B, Stellwagen NC. DNA persistence length revisited. Biopolymers;61(4):261–275. doi: 10.1002/bip.10151.

12. Kimura K, Hirano T. ATP-Dependent Positive Supercoiling of DNA by 13S Condensin: A Biochemical Implication for Chromosome Condensation. Cell. 1997;90(4):625–634. doi: https://doi.org/10.1016/S0092-8674(00)80524-3.

13. Doi M, Edwards SF. The Theory of Polymer Dynamics. OXFORD SCIENCE PUBLICATIONS; 1986.

14. Walther N, Hossain MJ, Politi AZ, Koch B, Kueblbeck M, degrd Fougner c, et al. A quantitative map of human Condensins provides new insights into mitotic chromosome architecture. Journal of Cell Biology. 2018;217(7):2309–2328. doi: 10.1083/jcb.201801048.

